# Functional Enrichment Analysis of Deregulated Long Non-Coding RNAs in Cancer Based on their Genomic Neighbors

**DOI:** 10.1101/2020.09.14.296921

**Authors:** Gulden Olgun, Oznur Tastan

**Affiliations:** Cancer Data Science Lab, National Cancer Institute, National Institute of Health, Sabanci University, Tuzla, Istanbul, Turkey; Faculty of Engineering and Natural Sciences, Sabanci University, Tuzla, Istanbul, Turkey

## Abstract

The dysregulation of long non-coding RNAs’ (lncRNAs) expressions has been implicated in cancer. Since most of the lncRNAs’ are not functionally characterized well, investigating the set of perturbed lncRNAs are is challenging. Existing methods that inspect lncRNAs functionally rely on the co-expressed coding genes, which are far better characterized functionally. LncRNAs can be known to act as transcriptional regulators; they may activate or repress the neighborhood’s coding genes on the genome. Based on this, in this work, we aim to analyze the deregulated lncRNAs in cancer by taking into account their ability to regulate nearby loci on the genome. We perform functional analysis on differentially expressed lncRNAs for 28 different cancers considering their adjacent coding genes. We identify that some deregulated lncRNAs are cancer-specific, but a substantial number of lncRNAs are shared across cancers. Next, we assess the similarities of the cancer types based on the functional enrichment of the deregulated lncRNA sets. We find some cancers are very similar in the functions and biological processes related to the deregulated lncRNAs. We observe that some of the cancers for which we find similarity can be linked through primary, metastatic site relations. We investigate the similarity of enriched functional terms for the deregulated lncRNAs and the mRNAs. We further assess the enriched functions’ similarity to the functions and processes that the known cancer driver genes take place. We believe that our methodology help to understand the impact of the lncRNAs in cancer functionally.

## 1 Introduction

Large scale profiling of the transcriptomes has revealed that the vast majority of the transcriptomes do not code for proteins [1, 2]. LncRNAs are the primary class of non-coding RNAs(ncRNAs) that are longer than 200 nucleotides. They undertake crucial roles in various biological mechanisms, including chromatin modification, transcriptional/post-transcriptional gene regulation [3]. Increasing evidence suggests that lncRNAs are key regulators in different cellular processes, and they can be used for therapeutic purposes. Dysregulation of lncRNAs is implicated in various human complex diseases such as cancer [4, 5, 6]. Therefore, understanding the functional mechanisms of lncRNAs is essential. However, to date, only a small number of lncRNAs are functionally characterized, which constitutes a challenge when analyzing the set of deregulated lncRNAs.

Several computational methods have been proposed for the functional analysis of lncRNAs [7, 8, 9, 10, 11]. Most of the computational methods rely on an analysis of mRNAs that are co-expressed with the lncRNA(s) in question. The LncRNA2Function [8] and lncRNAtor [9], Zhao et al. [10] infer lncRNA functions through analysis of the coding genes that are co-expressed with the lncRNAs in question. Linc2GO generates functional annotation for long intergenic non-coding RNAs (lincRNAs) based on the competing endogenous RNA hypothesis [7]. Unlike the previous methods, FARNA infers a transcript’s function from the transcription factors and co-factors whose genes are co-expressed with the transcript controlled by this TFs and TcoFs [11].

Most of the lncRNAs are expressed at lower levels compared to coding genes and, those lncRNAs are unlikely to regulate distal genes in a direct manner [12]. Therefore, inferring the functional activity of lncRNAs only from the co-expressed mRNAs may limit the analysis. LncRNAs act as local regulators on the genome, and they regulate nearby genes [13, 14, 15, 16]. Also, expression levels of the lncRNAs are associated with the expression of the neighborhood coding genes. Consequently, for a given IncRNA, we use its local regulation ability and the information on closeby genes to infer enriched biological processes and molecular functions.

In this work, we present a methodology for the functional investigation of deregulated lncRNAs in cancer. We first identify the set of differentially expressed lncRNAs for 28 different cancers, as obtained from The Cancer Genome Atlas Project (TCGA). We annotate lncRNAs with their neighboring coding RNAs based on gene-centric regulatory regions of mRNAs. We perform functional enrichment analysis on these coding genes. We further assess the functional similarities among cancers based on the semantic similarity of the enriched functions. We investigate whether these functional similarities suggest relationships between the cancer types based on the known primary and metastasis site interactions. To compare these findings, we apply the same pipeline for cancer driver genes and differentially expressed mRNAs in TCGA, and we compare enriched GO-terms. We further explore the functional mechanisms of cis-acting super-enhancer lncRNAs known to regulate the close-by mRNAs.

## 2 Methods

The overall methodology is summarized in Figure 1. For each of the deregulated lncRNA, we find the neighbor coding gene set. We apply enrichment analysis on the union of the coding gene sets for each lncRNAs and infer the related functions and processes of the deregulated lncRNAs. Based on these functions, we further analyze the relationship between these cancers. Below, we elaborate on those steps.

**Figure 1:**
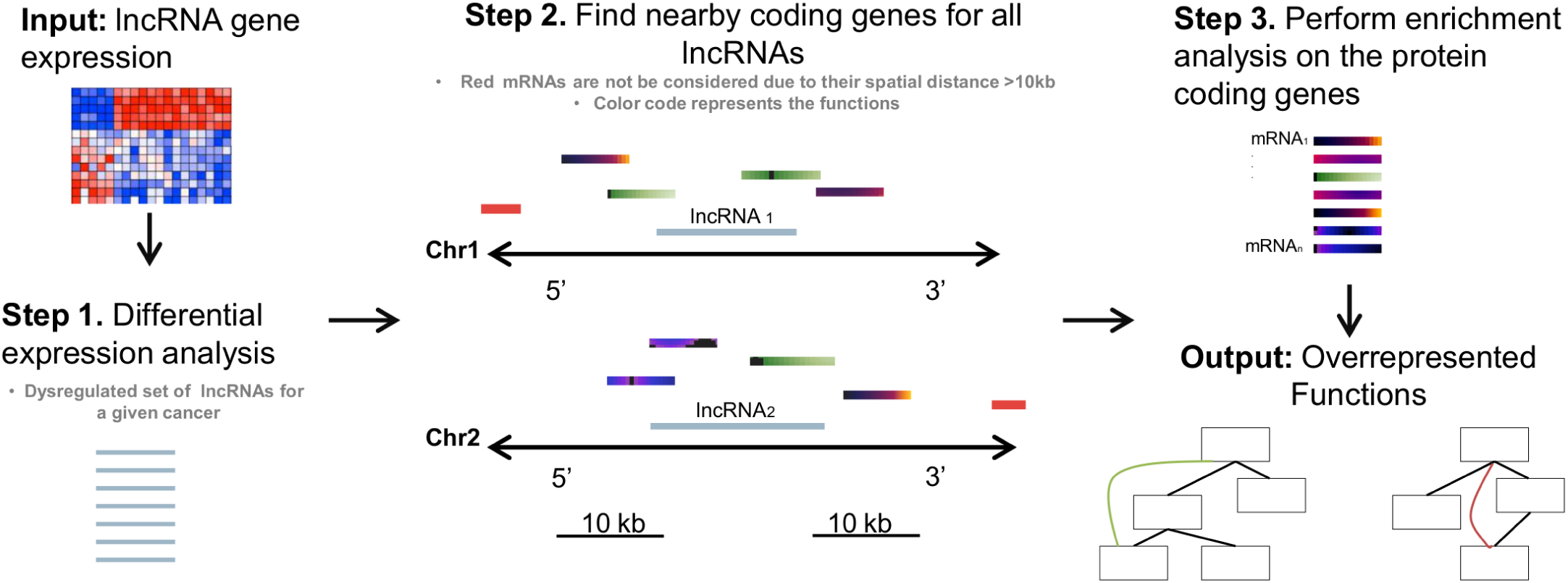
Overview of the methodology. Step1: Deregulated lncRNAs are determined by the differential expression analysis. Step 2: Within the 10 kb range from upstream and downstream of the lncRNA gene, lncRNA is annotated with their neighborhood coding genes. Step 3: From the annotated neighborhood coding gene set, we perform enrichment analysis. The same methodology is applied for 28 different cancer. Annotated coding genes are represented with different colors to reflect their functional variability. Also, mRNAs that are 10 kb far away from the lncRNA are illustrated with red color.

### 2.1 Data Sources and Data Processing

Illumina HiSeq RNA-Seq gene expression data for 28 different primary cancer sites are collected from GDC Data Portal (July 15^th^ 2017). We retrieve lncRNAs in RNASeq data from GENCODE v26 [17], which provides comprehensive gene annotation of lncRNA genes in the human transcriptome. In total, 15, 513 lncRNA genes are identified and considered in this study. Also, the remaining genes in RNA-Seq data are utilized for the mRNA validation.

Information on metastatic sites of each cancer is retrieved from MSK-IMPACT Clinical Sequencing Cohort [18] (downloaded on April 10^nd^ 2018). We utilize the cancer driver gene set as obtained from the IntOGen database [19]. Driver genes are collected on March 23^rd^ 2018. The cis-acting differentially expressed super-enhancer associated lncRNAs for different cancers are retrieved from the SELER database [20] (June 23^th^ 2019).

In our analyses, we use the fragments per kilobase of transcript per million mapped reads upper quartile normalization (FPKM-UQ) data as provided by TCGA. We filter out the lncRNA genes with an expression value of less than 0.05 in more than 20% of the samples. Expression values are added with a constant 1 to deal with the 0 gene expression values, and data were log 2 transformed. We eliminate the RNAs that do not vary across samples; we filter those with median absolute deviation (MAD), Equation 1, below 0.5.

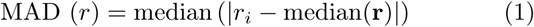

To identify the significantly deregulated lncRNAs, we use linear model fitting followed by the empirical Bayes t-test implemented in the Bioconductor limma package [21]. The significance level is set to 0.05.

### 2.2 Annotation and Functional Enrichment Analysis of lncRNAs

To get a functional insight on each differentially expressed lncRNA set for different cancer types, we conduct a functional enrichment analysis based on the neighboring genes. For this, we use our previously proposed method GLANET [22]. GLANET takes a genomic interval set defined by their genomic coordinates as input, in this case, the intervals being the lncRNAs. It then performs an enrichment analysis by calculating the overlap of the input with the genes in a functional gene-set. In the case of GO enrichment, the functional gene-set includes all genes annotated with a particular GO-term. When analyzing the pathway, it includes all genes in that pathway. Below we detail the two steps of this enrichment analysis.

#### 2.2.1 Finding Nearby Genes of lncRNAs based on Different Coding Genic Regions

GLANET also allows the user to define what genic regions of the functional gene-set should be considered in the analysis. For example, if the exon option is selected, only the exons of the genes in the functional gene set are considered when finding the overlap with the input set. These gene-centric regulatory regions define two different gene sets for enrichment, exon based, and regulation based gene sets. The exon based gene set comprises genes in the exon region. In contrast, the regulation based gene set contains introns and the four different gene-centric regulatory regions, namely 5p1, 5p2, 3p1, and 3p2, of these genes. The genomic intervals and the gene sets used in this analysis are illustrated in Figure 2 (A). Proximal regions, 5*p*1, and 5*p*2 are defined as the regions 0 − 2 kb and 2 − 10 kb upstream of the first exon of the gene, respectively. Similarly, 3*p*1 and 3*p*2 refer to the regions 0 − 2 kb and 2 − 10 kb downstream of the last exon of the gene, respectively.

**Figure 2:**
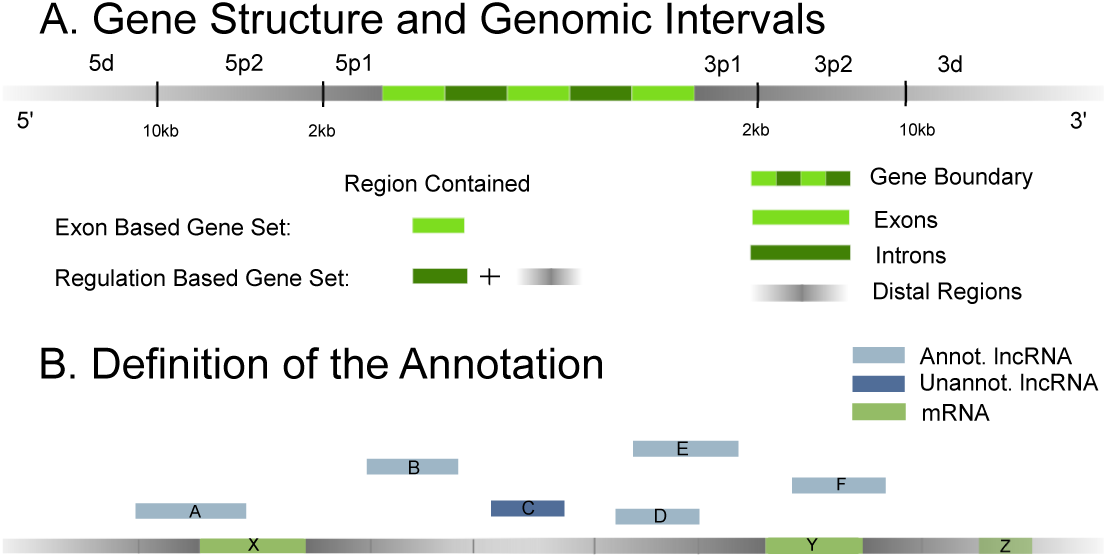
A. Illustration of the gene intervals and gene sets that are covered in this analysis. Distal regions are scaled down to fit the figure. B. The graphical representation of how a set of lncRNAs are annotated based on neighborhood coding genes. A lncRNA that falls into any of the genomic intervals of mRNA is annotated with the corresponding gene and this mRNA is added to the related genomic gene set. For a given physical map, IncRNA A is annotated with mRNA X in the exon and regulation based gene set. lncRNA F, on the other hand, is annotated with mRNA Y for the exon based gene set and with mRNA Y and Z on the regulation gene set.. lncRNA B is only annotated with mRNA X in the regulation based gene set. LncRNA D and E are annotated with mRNA Y in a regulation based gene set. Finally, lncRNA C falls into the 3D/5D genomic intervals of mRNA Y, and as a result, it is not annotated with mRNAs in any gene set. Thus, for enrichment analysis, the exon based gene set consists of mRNA X and Y. Similarly, the regulation based gene set contains mRNA X, Y, and Z.

When finding a nearby gene of a lncRNA in the input list, the overlapping mRNA gene list is curated based on whether an exon-based or regulation-based analysis option is selected. Once the overlapping mRNA gene set is defined, the functional enrichment analysis is conducted using that set of coding genes. This annotation idea is summarized in Figure 2 (B).

#### 2.2.2 Functional Enrichment Analysis of lncRNAs

Subsequently, the functional enrichment step assesses the significance of the observed overlap with the input lncRNA set with that of the functional gene set. For quantifying the overlaps, we consider the length of the overlapping region. The sampling-based enrichment tests are conducted with 10, 000 samplings. The random intervals are generated by matching the input’s chromosome, length, mappability, and GC content of the input interval set (for details see [22]). The GO-terms and KEGG pathways that achieve significance with Benjamini-Hochberg correction with *FDR <* 0.05 are considered as significantly enriched.

### 2.3 GO Term Semantic Similarity Evaluation

To asses how cancers show similarity based on the functions of the deregulated lncRNAs, we calculate the GO semantic similarity between the enriched GO term sets of two cancers. Let the enriched GO-term set for the first cancer be *GO*_*x*_ = {*go*_*x*1_, *go*_*x*2_, *go*_*x*3_,.., *go*_*xk*_} and genes in the second cancer are enriched with the GO-terms, *GO*_*y*_ = {*go*_*y*1_, *go*_*y*2_, *go*_*y*3_, …, *go*_*yl*_}. The similarity score between the two GO-term sets is calculated with the graph-based Wang method [23], Equation 2. When calculating the semantic similarity between the two GO-terms, the Wang method considers the position of the GO-terms in the GO hierarchy and their relations with their ancestor terms.

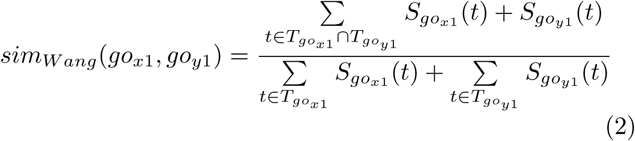

where 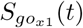 is the contribution of a GO-term *t* to the semantics of GO-term *go*_*x*1_ and 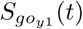 is the contribution of a GO-term *t* to the semantics of GO-term *go*_*y*1_. 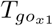 and 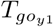 denote the terms related to the terms in the GO graph. To calculate the semantic similarity among sets of GO-terms we use the Best-Match average strategy (BMA) method which computes the average of all maximum similarities on each row and column of the pairwise GO-term similarity table as follows:

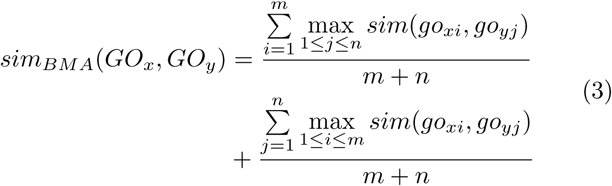

### 2.4 Clustering Cancers

We cluster the cancers using hierarchical clustering with the average linkage method. The cancers’ similarities are based on the semantic GO term similarity calculated over the enriched GO terms in that cancer. The optimal cluster size is determined using the Dunn index [24]. Dunn index measures the compactness and separation of the clusters by calculating the ratio between the smallest distance between observations not in the same cluster to the largest intra-cluster distance.

## 3 Results

### 3.1 Overview of Gene Ontology Enrichment Analysis of the lncRNAs

We conduct the analysis summarized in Figure 1 to analyze the deregulated lncRNAs in cancers. In this analysis, the lncRNA enrichment analysis is based on the nearby genes on the genome. The exon-based and regulation-based analyses are carried out to understand the effect of different regions. In the first step, we identified the significantly deregulated lncRNAs in cancer. The number of lncRNAs for each cancer that are curated after differential expression analysis is provided in Figure 3 (*p*-value *<* 0.05). The same figure also depicts the number of tumor patients that are covered in this analysis for each cancer type. We find that 1, 186 differentially expressed lncRNAs are common in all cancer types. This may indicate that there common mechanisms in cancer that are perturbed, in which lncRNAs are involved.

**Figure 3:**
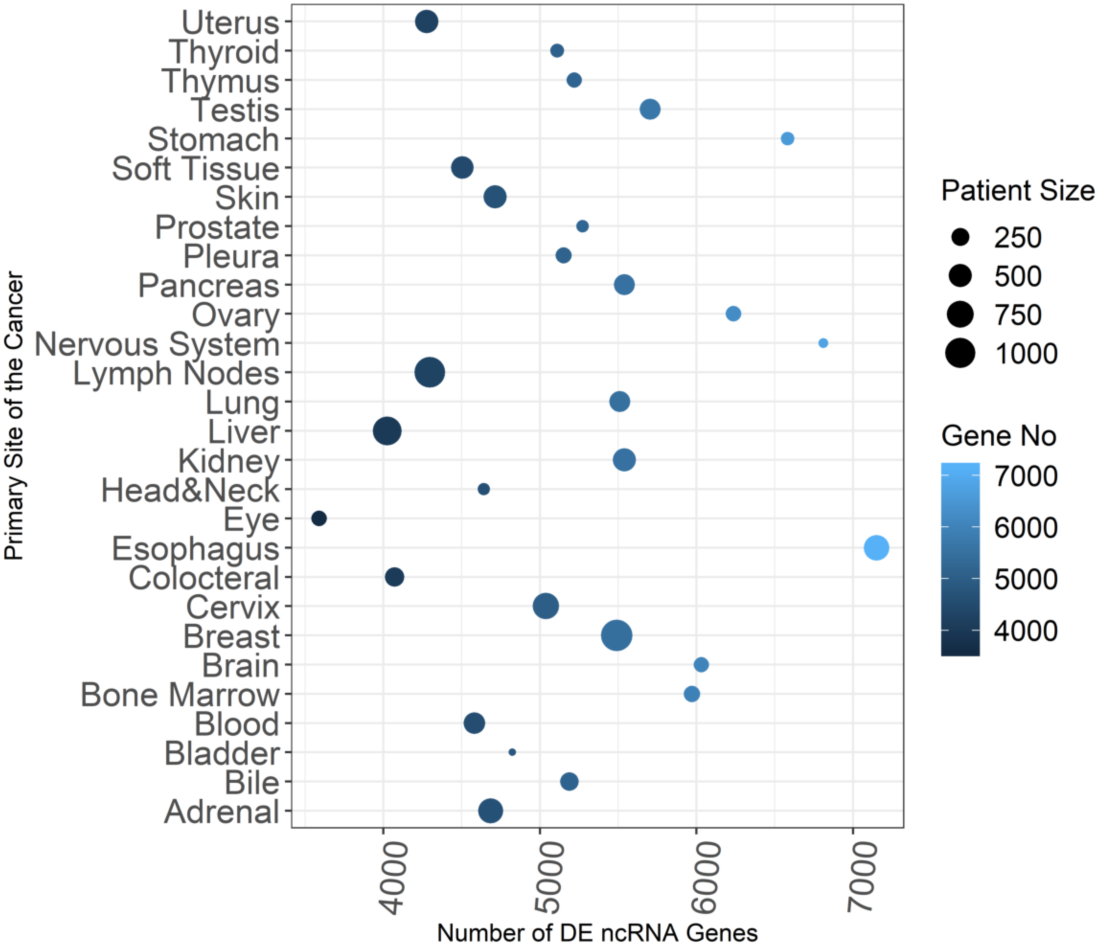
Dot plot for the number of tumor patients and the number of differentially expressed lncRNAs in each cancer. The color code of the dots represents the number of differentially expressed lncRNA genes and the size of dots shows the number of tumor patients for cancer.

We find that some enriched GO-terms are common in all cancers. Among these common GO-terms, most of them are related to signaling, which is associated with cancer [25]. Exon-based analysis of the lncRNAs results with enriched terms related to the transcription regulation, and biosynthetic related biological processes for all cancer types. In the regulation-based analysis, most of the enriched biological processes are related to regulation, morphogenesis, development, metanephric except nervous, ovarian, pancreas, esophagus, and stomach cancers. Increased expression in fatty acid translocase, fatty acid transport protein family, and plasma membrane fatty acid-binding proteins are previously reported in cancer, including ovarian cancer [26]. Most of the enriched GO-terms for regulation based analysis of the lncRNAs are related to long-chain fatty acids in ovarian cancer. Studies reveal that increased lipogenesis based on a fatty acid desaturase observed in ovarian cancer [27]. In the pancreas and esophagus cancer, most of the enriched GO-terms for regulation based analysis of the lncRNAs are related to the secretion. This is expected due to the secretion ability of the pancreas and esophagus.

### 3.2 LncRNA based Functional Similarity of Cancers

We identified groups of cancers that bear similarities in terms of the functions of the deregulated lncRNA functions. We clustered the cancers based on the similarity of the enriched functions in the deregulated cancers and repeated this analysis for exon-based and regulation-based. The heatmap representations of the semantic similarity scores between cancers based on GO molecular function and biological processes are shown in Figure 4. In this figure, clusters are demonstrated with black squares, and the optimal cluster size is determined with the Dunn index (see Section 2). We observe exon based analysis reveals have more defined clusters than regulation based analysis

**Figure 4:**
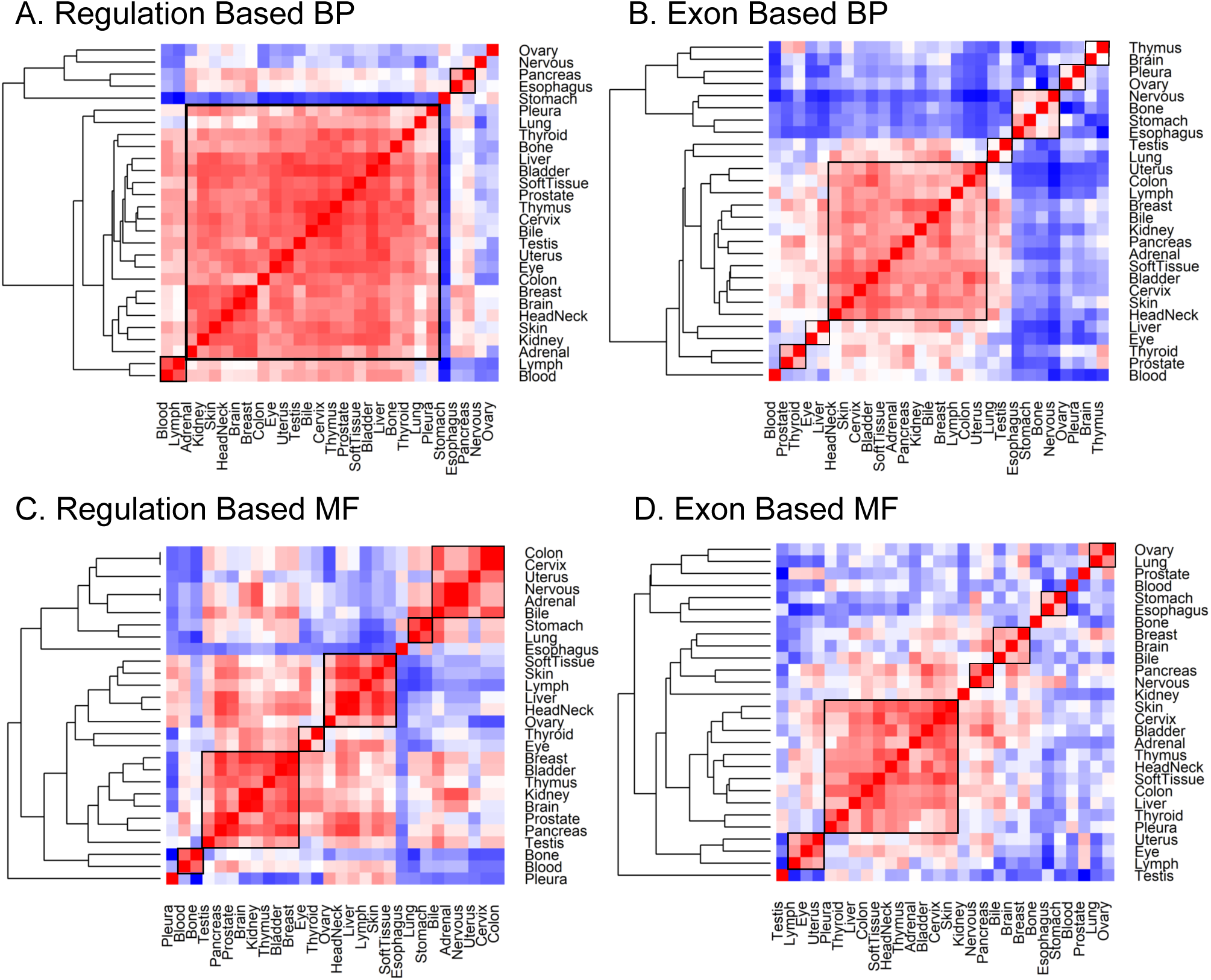
The functional similarity score between each cancer pairs is represented. Biological process analyses (A-B) are demonstrated in the first row and molecular functional analyses (C-D) are in the second row. Colors are scaled between blue (low) to red (high) and white color represents the median value. Color key for regulation based heatmap is [0.2-1] and for exon based is [0.4-1]. Clusters are highlighted with black squares. Row and column labels describe the primary site of the cancers.

We observe some similarities across cancers. In the regulation based analysis, we detect the Diffuse Large B-cell Lymphoma (DLBL) and Acute Myeloid Leukemia (AML) emerge as very similar, Figure 4 (A). In this figure, primary site *blood* represents AML, and primary site *lymph* represents DLBL. DLBL is the most common subtype of non-Hodgkin’s lymphoma that affects B-cells, one type of white blood cell responsible for producing antibodies [28]. Acute Myeloid Leukemia is abnormally increased in the number of the myeloid line of blood cells in the bone marrow. Interestingly, this similarity is detected only in the regulation-based analysis. Biological process GO-terms that enrich only in both AML and DLBL is listed in Table 1. It was previously reported that *K63-linked ubiquitination* is linked to the AML and DLBL [29]. Moreover, biosynthetic processes allow a cell to develop macromolecules to cause abnormal cell division, and tumor growth is essential for cancer [30]. It is known that biosynthetic processes inhibit AML and DLBL, but the specific mechanisms still are not clear yet [31].

**Table 1:**
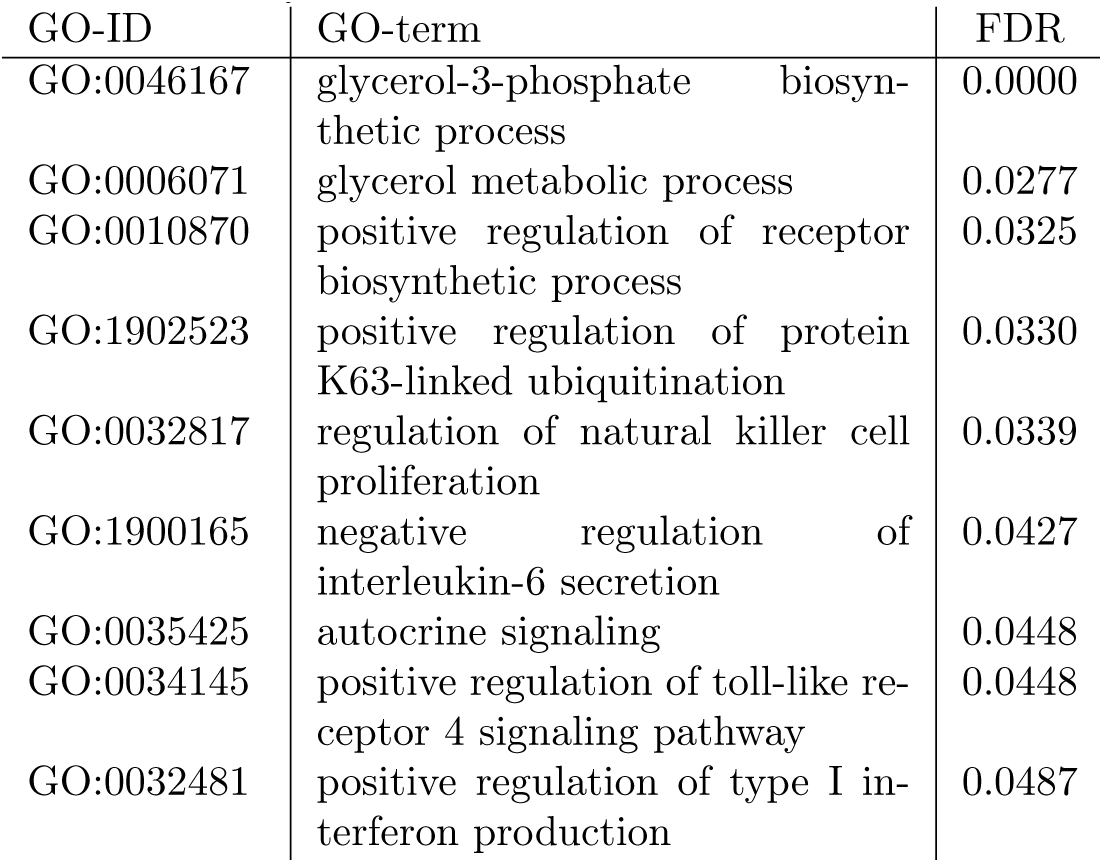
The list of biological process GO-terms that only enriched in both Acute Myeloid Leukemia and Diffuse Large B-cell Lymphoma. The last column shows the maximum FDR value that AML and DLBL have for the corresponding GO-terms when they enriched.

| GO-ID | GO-term | FDR |
| --- | --- | --- |
| GO:0046167 | glycerol-3-phosphate biosynthetic process | 0.0000 |
| GO:0006071 | glycerol metabolic process | 0.0277 |
| GO:0010870 | positive regulation of receptor biosynthetic process | 0.0325 |
| GO:1902523 | positive regulation of protein K63-linked ubiquitination | 0.0330 |
| GO:0032817 | regulation of natural killer cell proliferation | 0.0339 |
| GO:1900165 | negative regulation of interleukin-6 secretion | 0.0427 |
| GO:0035425 | autocrine signaling | 0.0448 |
| GO:0034145 | positive regulation of toll-like receptor 4 signaling pathway | 0.0448 |
| GO:0032481 | positive regulation of type I interferon production | 0.0487 |

We also observe an interesting similarity of the stomach and esophageal carcinoma across a molecular function. Stomach adenocarcinoma occurs in the mucus-producing cells in the stomach, whereas esophageal carcinoma develops in the esophagus’s glands, fibromuscular tube that connects the throats to the stomach. In the exon-based analysis, the molecular function GO-term *acetyltransferase activator activity (GO:0010698)* was found enriched only in the stomach and esophageal carcinoma. Recent studies show the link between acetyltransferases and gastric cancer [32, 33]. Thus, lncRNAs may be the local regulators of these coding genes that participate in these activities, and lncRNAs may indirectly promote cancer. Additionally, we detect *fibroblast growth factor binding (GO:0017134)* and *C-X3-C chemokine binding (GO:0019960)* molecular functions which are important for both stomach adenocarcinoma and esophageal carcinoma [34, 35].

### 3.3 The Primary-Metastatic Site Relations of the Clustered Cancers

We analyze the cancer cluster to explore whether these cancers are related due to the metastatic sites. We subset the MKS-IMPACT data for the 28 TCGA tissue sites. MSK-IMPACT data lists sub-cancer types. The primary and metastatic site of each patient in MSK-IMPACT data is matched with the corresponding TCGA site. Figure 5 shows the directed graph of cancers. In this network, each node corresponds to a cancer site. An edge indicates from the primary site to the metastatic site if it is observed. Every primary tumor except the blood cancer metastases to another tissue.

**Figure 5:**
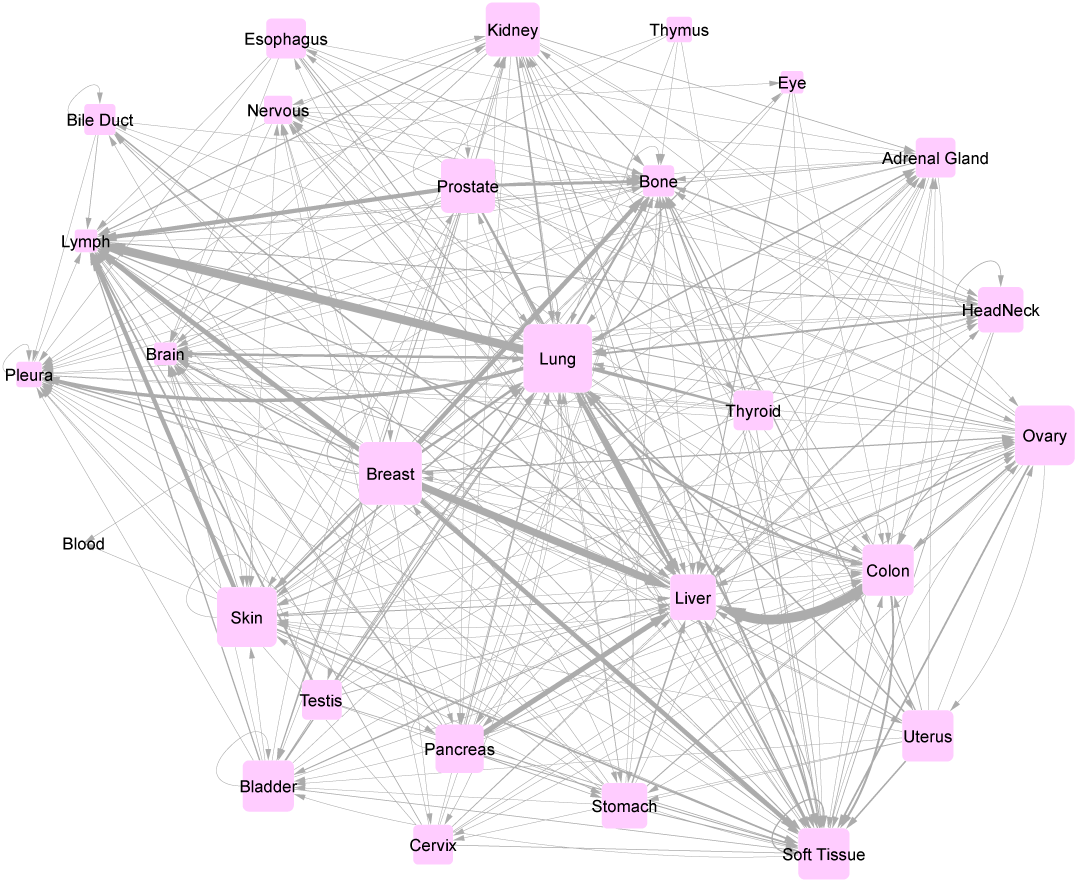
The metastasis network based on MSK-IMPACT data [18]. Each node corresponds to a cancer site. The edges link the primary sites to the metastasis sites. The size of a node depends on the out-degree of the node, the number of outward-directed edges from that node. The weights of edges are the number of occurrences of the related metastasis for the corresponding node.

Uveal melanoma is the most common ocular malignancy in the iris, ciliary body, or choroid [36]. Metastasis is observed almost in 50% of patients within 6-12 months of survival [36]. Most of the metastasis is formed in the liver [37]. Our findings in exon based analysis, Figure 4 (B), shows that primary site liver and eye have similar functional patterns in the biological process. When examining the metastasis in the liver and eye, 21 cancer patients, whose primary sites are eye, have liver metastasis, in MSK-IMPACT data (Figure 5).

Colon cancer is the third leading cause of cancer-related deaths in 2018 [38]. The liver is the most common site of the colon cancer metastasis, more than 50% of the colon cancers [39] due to the most of the intestinal mesenteric drainage joins the hepatic portal venous system. The most common metastasis in MSK-IMPACT data is the primary site colon causing liver metastasis. 297 patients whose primary site is colon cancer suffer from liver metastasis, Figure 5. Interestingly, colon and liver cancers have similar functional pattern and cluster together in both regulation based lncRNA biological process, and exon based lncRNA molecular function analysis.

### 3.4 Pathway Analysis

We include the pathway analysis for a set of lncRNAs that are derived from their neighborhood genes. Pathway enrichment analysis for lncRNAs derived from their neighborhood gene is conducted with KEGG pathways using GLANET. We detect few enriched KEGG pathways for regulation and exon based gene sets. The regulation based analysis for prostate cancer is enriched with *pancreatic secretion* (*hsa04972, FDR* = 0.026) pathway. Previous studies do not provide evidence on the link to this. However, we may point an indirect link between prostate cancer and pancreatic secretion. Serine peptidase inhibitor Kazal Type 1, SPINK1 gene, is secreted from pancreatic acinar cells into pancreatic juice. Its role is to prevent trypsin-catalyzed premature activation of zymogens within the pancreas and the pancreatic duct [40]. Moreover, it has been associated with prostate cancer [41]. Thus, prostate cancer and *pancreatic secretion (hsa04972)* pathway may be related, and lncRNAs of prostate cancer may participate in the transcription or splicing in tumorigenesis.

*Phenylalanine metabolism (hsa00360, FDR <* 0.05) pathway is enriched in the prostate, liver, breast, brain, and bile duct cancer for exon based lncRNAs. Phenylalanine is the essential amino acid that naturally found in breast milk and also cheese, eggs, and meat. It is utilized in the dopamine and norepinephrine neurotransmitters. Thus, it is intensively presented at the human brain and plasma. Phenylalanine is the source of the artificial sweetener aspartame [42], which is claimed to caused brain cancer [43, 44]. However, we could not find a reported relation between phenylalanine metabolism pathway and the other cancers, where it was found to be enriched.

### 3.5 Comparing the Enriched Terms in the Deregulated lncRNA set With the Enriched Terms in the Coding Genes

We conduct two analyses in the first analysis; we focus on the deregulated coding mRNAs. In the second one, we compare the enriched functions and processes of cancer driver genes in IntOGen [19]. We elaborate the details of how we perform these analyses in the following subsections:

#### 3.5.1 Identifying Functional Relevance between lncRNAs and mRNAs

We examine the functional relevance between a set of deregulated lncRNAs and a set of deregulated mRNAs for each cancer and state whether they participate in similar functions. For this purpose, we employ the same pipeline to the mRNA genes in TCGA RNASeq data. We also use fragments per kilobase of transcript per million mapped reads upper quartile normalization (FPKM-UQ). The same pre-processing step is applied, and differential expression analysis is performed with the limma package for *p* − *value <* 0.05 and *FDR <* 0.05 cut-offs. A set of differentially expressed mRNA for each cancer is annotated with biological process GO-terms, and GO-term enrichment analysis is performed with GLANET [22].

We compare enriched GO-terms obtained for the exon based mRNA enrichment analysis and the exon based lncRNA enrichment analysis. The functional semantic similarity between two sets for each cancer is listed in Table 2. We observe a high functional semantic similarity score (*>* 0.87) between lncRNA and mRNA. It indicates that the deregulated lncRNAs may contribute to similar functions with the deregulated mRNAs through regulating their nearby genes.

**Table 2:**
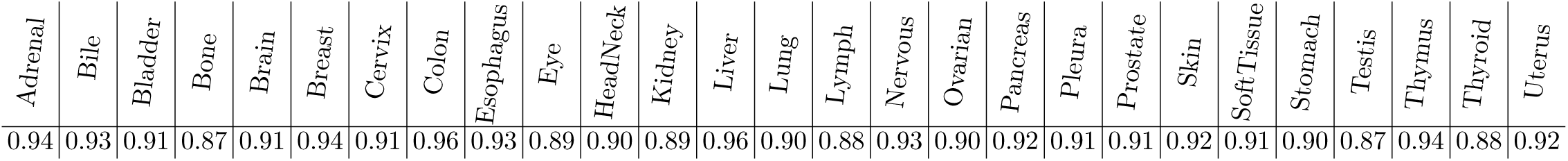
The functional semantic similarity scores between the differentially expressed mRNA and lncRNA gene sets calculated for each cancer.

#### 3.5.2 Comparing the Enriched Functions and Process with the Cancer Driver Genes’ Function and Processes

This analysis aims to understand whether the deregulated lncRNAs share similar functional mechanisms as the driver genes for a given cancer. In this analysis, we collect the driver genes from IntOGen database [19]. We carry out the enrichment tests with clusterProfiler [45] for each cancer. *p*-value and *FDR* cut-offs for this analysis are set to 0.05. The semantic similarity scores are evaluated between enriched GO-terms for cancer drivers and the enriched terms for the deregulated lncRNAs (see *Materials and Methods* section). The semantic scores are listed in Table 3. The driver gene pool is small and functional heterogeneity may be limited. Thus, we obtain low semantic functional similarity scores between driver genes and both exon and regulation based analysis of the lncRNAs.

**Table 3:** The semantic similarity scores between enriched GO-terms in the caner driver gene set and the ones obtained through analysis of deregulated lncRNAs for corresponding cancer. Also, empirical p-values for the permutation test are provided.

|  | BP |  |  |  | MF |  |  |  |
| --- | --- | --- | --- | --- | --- | --- | --- | --- |
|  | Similarity Score |  | Empirical p-value |  | Similarity Score |  | Empirical p-value |  |
|  | Exon | Reg. | Exon | Reg. | Exon | Reg. | Exon | Reg. |
| Stomach | 0.444 | 0.188 | 0.050 | 0.137 | 0.492 | 0.343 | 0.054 | 0.069 |
| Adrenal | 0.392 | 0.391 | 0.007 | 0.007 | 0.414 | 0.214 | 0.015 | 0.018 |
| Breast | 0.410 | 0.353 | 0.060 | 0.104 | 0.404 | 0.274 | 0.040 | 0.061 |
| Bladder | 0.440 | 0.400 | 0.020 | 0.025 | 0.432 | 0.284 | 0.034 | 0.020 |
| Blood | 0.450 | 0.449 | 0.025 | 0.027 | 0.401 | 0.485 | 0.025 | 0.022 |
| Bone | 0.425 | 0.390 | 0.025 | 0.024 | 0.380 | 0.370 | 0.020 | 0.021 |
| Brain | 0.356 | 0.340 | 0.030 | 0.049 | 0.444 | 0.238 | 0.044 | 0.053 |
| Colon | 0.433 | 0.396 | 0.024 | 0.018 | 0.369 | 0.177 | 0.017 | 0.042 |
| Esophagus | 0.460 | 0.340 | 0.022 | 0.025 | 0.360 | 0.306 | 0.026 | 0.034 |
| Head&Neck | 0.440 | 0.345 | 0.035 | 0.082 | 0.384 | 0.348 | 0.056 | 0.052 |
| Kidney | 0.426 | 0.362 | 0.029 | 0.037 | 0.378 | 0.254 | 0.028 | 0.040 |
| Liver | 0.416 | 0.422 | 0.013 | 0.011 | 0.416 | 0.335 | 0.022 | 0.020 |
| Lung | 0.419 | 0.335 | 0.240 | 0.756 | 0.408 | 0.167 | 0.047 | 0.249 |
| Ovary | 0.363 | 0.231 | 0.026 | 0.028 | 0.443 | 0.301 | 0.021 | 0.021 |
| Pancreas | 0.363 | 0.291 | 0.011 | 0.012 | 0.463 | 0.334 | 0.011 | 0.009 |
| Prostate | 0.400 | 0.448 | 0.018 | 0.011 | 0.526 | 0.442 | 0.018 | 0.019 |
| Skin | 0.405 | 0.337 | 0.791 | 0.849 | 0.404 | 0.353 | 0.117 | 0.376 |
| Thyroid | 0.451 | 0.418 | 0.011 | 0.009 | 0.449 | 0.272 | 0.016 | 0.011 |
| Uterus | 0.426 | 0.393 | 0.045 | 0.102 | 0.442 | 0.203 | 0.041 | 0.081 |

We further define a null model to assess how interesting is the semantic similarity scores. We use a random sampling-based approach. We randomly select driver genes from the pool of cancer driver genes that includes all cancers, not only particular cancer we are interested in. We draw on these randomized gene set to get an empirical null distribution of the test statistic. We calculate the empirical *p* − *value* on this null distribution by measuring the number of cases where similarity scores of the randomly selected driver genes are greater than the observed GO-term similarity score. 1, 000 random samplings are used. Note that since many cancers share drivers, it is a stringent test to pass. The *p* − *values* of these tests are provided in Table 3. In some cases, significant results are attained in most cases, providing evidence that the deregulated lncRNAs may be taking a role in the same processes and functions as the cancer driver genes.

When we compare the enrichment results, we discover that our lncRNA annotation and enrichment pipeline helps identify the critical functions that would have been missed solely with a mRNA based analysis. For example, in the breast cancer analysis, our method identifies phosphatidylserine biosynthesis-related biological functions, whereas an analysis that is limited to the mRNAs fail to detect these functions. Phosphatidylserines have been reported to participate in cell cycle signaling, especially in apoptosis [46, 47]. And differences of the exposure of phosphatidylserine in the cancer cells but in the healthy cells have been reported, which is utilized for therapeutic purposes [47].

### 3.6 Inferring Functional Mechanisms for the Super-enhancer Associated lncRNAs

Enhancers are cis-regulatory DNA elements that increase the transcription of a gene. Super-enhancers are groups of putative enhancers that are spatially proximal to the mediator binding sites [48]. They regulate gene expression by increasing the gene transcription [20]. Disease-related variations occur in the super-enhancer regions. Therefore, super-enhancers are associated mostly with diseases, including cancer, and they can be used as biomarkers [49]. The recent study reveals cis-acting super-enhancer lncRNAs are transcripted from the super-enhancers and regulate closeby genes. Thus, we analyze these genes’ functional mechanisms based on their neighborhood coding genes using their cis-regulation ability.

In this analysis, we collect the differentially expressed cis-acting lncRNAs from the SELER database. We only consider the genes that more than 20% of the samples expressed higher than the 0.05 expression value and varied across samples (see Methods). The final set contains 65 lncRNA for bladder cancer, 49 lncRNA for breast cancer, 179 lncRNA for kidney cancer, 97 lncRNA for lung cancer, 47 for prostate cancer, 42, and 50 lncRNAs for stomach and thyroid cancer, respectively. We apply the same methodology for the cis-acting super-enhancer lncRNAs. We identify the set of coding genes proximal to those cis-acting super-enhancer lncRNAs, and we perform gene ontology and pathway enrichment analysis on these coding genes.

We observe that regulation based related GO-terms such as positive regulation of telomerase activity, negative regulation of transcription by competitive promoter binding are enriched for all cancers. Interestingly, excluding stomach cancer, comprise the cell death-related GO-terms such as negative/positive regulation of programmed cell death, autophagic cell death, engulfment of target by autophagosome, cell death, and other apoptosis-related GO-terms are also enriched. This may indicate their pathogenesis property.

### 3.7 Comparison with the Other lncRNA Functional Analysis Methods

To evaluate our pipeline, we compare our results with the existing lncRNA functional analysis methods. The details of these analyses are explained in the following sections:

#### 3.7.1 Comparison for the Gene Ontology Enrichment Results

As a case study, we limit our studies to breast cancer and use only differentially expressed HOTAIR, H19, and MEG3 lncRNA genes. The purpose of selecting those genes is they are experimentally proved to be associated with tumorigenesis of breast cancer [50, 51]. To implement this analysis, we only perform enrichment analysis of those lncRNAs with GLANET [22] and compare the results with the other methods. Co-LncRNA, FARNA, Linc2GO, LncRNA2Function, and lncRNAtor methods are considered. However; we utilize only Co-LncRNA and FARNA [10, 11] for the comparative analysis. Linc2GO [7] and LncRNA2Function [8] are no longer accessible. Moreover, lncRNAtor [9] does not support breast cancer data. FARNA allows inputting only single lncRNA. Thus, enrichment results are compared on an individual basis in FARNA, even though our pipeline is designed for a set of lncRNAs. Our analysis and FARNA, mutually, identify negative regulation of *cell proliferation (GO:0008285)* GO-term for H19 and HOTAIR. However, mutual enrichment result for lncRNA MEG3 is not detected. Co-LncRNA investigates the combinatorial effects of the lncRNAs. Therefore, it allows only to use a maximum of three lncRNAs for functional enrichment analysis. To perform enrichment analysis in Co-LncRNA, we use linear Regression and all *p*-value cut-offs set to 0.05. *Negative regulation of cell proliferation (GO:0008285), positive regulation of transcription, DNA-dependent (GO:0045893), negative regulation of transcription, DNA-dependent (GO:0045892)* biological process GO-terms are detected mutually in Co-LncRNA and our proposed method. Moreover, *transcription coactivator activity* and *RNA polymerase II core promoter proximal region sequence-specific DNA binding* GO-terms are found commonly for molecular function GO-terms.

Our method finds the *gonadotropin secretion (GO:0032274)* function as enriched in both exon-based and regulation-based analyses. Interestingly, none of the compared methods identifies this function for the same input. Gonadotropin hormone-releasing hormones are synthesized by the anterior pituitary and also produced in pregnancy. They are extensively used in the treatment of hormone-dependent diseases such as breast cancer [52].

#### 3.7.2 Comparison for the Pathway Enrichment Results

We also compare our results with LncRNAs2Pathways [53]. LncRNAs2Pathways is an R-based tool under a LncPath package name. It identifies the pathways affected by a set of lncRNAs, based on a Kolmogorov-Smirnov like statistical measure weighted by the network propagation score. It considers the correlation of expression between the lncRNAs and the coding mRNAs. To perform LncRNAs2Pathways analysis, we built the coding-non-coding gene correlation (CNC) network where nodes are protein-coding and ncRNA genes, edges are interactions between co-expressed protein-coding and ncRNA gene pairs. To construct a CNC network, we apply the same pre-processing step, and differential expression analysis for expression profiles of the lncRNA and mRNA genes (See Methods section). Pearson correlation coefficient is calculated for each differentially expressed lncRNA-mRNA pair. Co-expression patterns between lncRNA and mRNA are detected; if their coefficient value ranked in the top or bottom of the 0.1% and *p*-value *<* 0.01. We apply stringent cut-offs to identify highly correlated pairs. In LncRNAs2Pathways analysis, we use brain cancer as a case study. Given the CNC network, the *Ribosome pathway* is enriched in brain cancer (*FDR <* 0.05) with LncRNAs2Pathways. We could not find a direct relationship between the brain and the ribosome pathway, even though our proposed method finds related pathways.

## 4 Conclusion

Existing methods that infer lncRNA function relies on the functional analysis of co-expressed coding genes. In this work, we present an alternative strategy where we functionally analyze the deregulated lncRNAs in different cancers through coding genes that are nearby on the genome. We also elucidate the functional relationship between cancers based on the enriched functions of the deregulated lncRNAs.

Our analysis depends on the completeness of the functional annotations of the coding genes. Some coding genes are functionally characterized better than the others. Improvements in the GO-term annotations will enhance our analysis and make our analysis more robust. Similarly, as the lncRNAs are functionally characterized further, our understanding of how they influence the cancer mechanisms will be clearer.

